# Modeling implanted metals in electrical stimulation applications

**DOI:** 10.1101/2021.12.04.471211

**Authors:** Borja Mercadal, Ricardo Salvador, Maria Chiara Biagi, Fabrice Bartolomei, Fabrice Wendling, Giulio Ruffini

## Abstract

**Background:** Metal implants impact the dosimetry assessment in electrical stimulation techniques. Therefore, they need to be included in numerical models. While currents in the body are ionic, metals only allow electron transport. In fact, charge transfer between tissues and metals requires electric fields to drive the electrochemical reactions at the interface. Thus, metal implants may act as insulators or as conductors depending on the scenario.

**Objective/Hypothesis:** The aim of this paper is to provide a theoretical argument that guides the choice of the correct representation of metal implants using purely electrical models but considering the electrochemical nature of the problem in the technology of interest.

**Methods:** We built a simple model of a metal implant exposed to a homogeneous electric field of various magnitudes to represent both weak (e.g., tDCS), medium (TMS) or strong field stimulation. The same geometry was solved using two different models: a purely electric one (with different conductivities for the implant), and an electrochemical one. As an example of application, we also modeled a transcranial electrical stimulation (tES) treatment in a realistic head model with a skull plate using a high and low conductivity value for the plate.

**Results:** Metal implants generally act as electric insulators when exposed to electric fields up to around 100 V/m (tES and TMS range) and they only resemble a perfect conductor for fields in the order of 1000 V/m and above. The results are independent of the implant’s metal, but they depend on its geometry.

**Conclusion(s):** Metal implants can be accurately represented by a simple electrical model of constant conductivity, but an incorrect model choice can lead to large errors in the dosimetry assessment. In particular, tES modeling with implants incorrectly treated as conductors can lead to errors of 50% in induced fields or more. Our results can be used as a guide to select the correct model in each scenario.

## Introduction

The presence of implanted metals in the body during electrical stimulation alters the current flow impacting the electric field magnitude and distribution. In all electrical stimulation modalities, this raises safety concerns [1] or, at the very least, it has an impact on the dosimetry assessment [2] or optimization [3]. In fact, while due to these concerns, in many transcranial electrical stimulation (tES) studies metal implants are included as part of exclusion criteria [4], in some populations, such as patients with epilepsy, this may severely limit recruitment numbers and translational impact (this population is susceptible to have deep brain stimulation implanted electrodes [5,6] or skull plates implanted after a craniotomy [7]). In brain stimulation techniques such as tES or transcranial magnetic stimulation (TMS), the electric field distribution in the brain is usually estimated using patient-specific models solved with the finite elements method (FEM) [8,9]. For an accurate estimation of the fields, it is therefore important to properly model metal implants if they are present.

Metal implants have electrical conductivity values several orders of magnitude higher than those of biological tissues. Thus, it may seem straightforward to model metals by associating them a very large conductivity or by simply assuming they act as perfect conductors. This approach is indeed very common in the literature [2,10]. However, while currents in the body are ionic in nature, metals only allow for electron transport. In fact, although metals are very good conductors, charge transfer between biological tissues and metals requires electrochemical reactions to happen at the interface [11], which require energy, i.e., presenting a voltage difference at the interface. This means that, depending on the interface voltage, implanted metals may act as insulators or as conductors.

In many applications, the electric field magnitude generated around metal implants is relatively low and, therefore, a low voltage gradient at the interface of an implant is produced. In this scenario metal implants are expected to act more like insulators than perfect conductors. This has already been suggested by other authors in the context of tES [12]. However, it seems general practice to model implanted metals as conductors regardless of the context. Indeed, with a quick search one can find many studies where metals are modeled as highly conductive in situations where they may have been better represented as insulators. To mention a few examples, non-active contacts in deep brain stimulation [13] or peripheral nerve stimulation [14], and orthopedic prostheses [15] have been modeled as highly conductive. This may be explained by the fact that, to the best of our knowledge, neither numerical studies nor theoretical arguments can be found in the electrical stimulation literature that would justify modeling metal implants differently than as highly conductive.

In the present study we aim at raising awareness on the issue described above and provide guidelines to correctly model passive implanted metals. For this purpose, we analyze the results of numerical models when the interface between a metal and tissue is modeled considering the electrochemical nature of the problem compared to the results of a purely electrical model. In addition, we simulate a realistic scenario of tES treatment on a subject with a skull plate to illustrate the importance of the correct model choice for the plate surface.

## Materials and methods

### Simple electrochemical box model

A simple model of a cylindrical implant exposed to a homogeneous electric field was built using the finite elements method (FEM) software platform COMSOL Multiphysics v5.3a (Stockholm, Sweden). The geometry of the model, depicted in Figure 1a, consisted of a cylinder (2 mm diameter and 10 mm length) embedded in a cube with 20 mm side. The geometry was meshed using COMSOL built in tools (see Figure 1b) resulting in a mesh of 411,469 first-order tetrahedral elements with an average element quality of 0.71.

**Figure 1:**
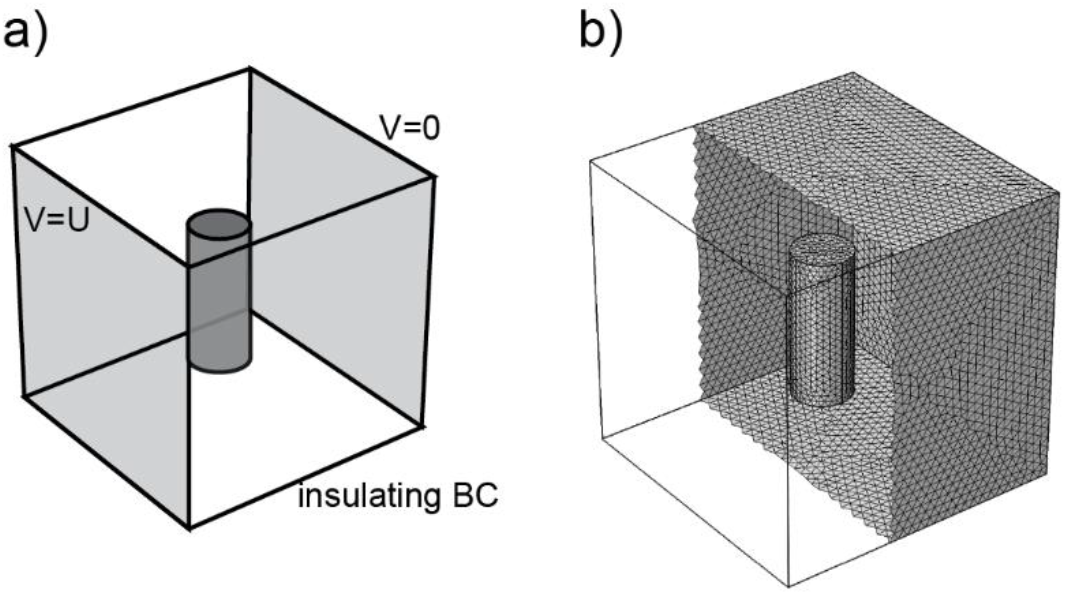
a) Schematic of the simple model of a cylindrical implant exposed to a homogeneous electric field. b) Tetrahedral mesh generated to solve the model using FEM.

The electric potential distribution was calculated by solving Laplace equation over the entire volume

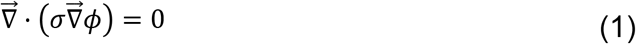

were *ϕ* is the voltage and *σ* is the electrical conductivity. To simulate a homogenous electric field, a voltage difference between two opposite sides was generated by defining a Dirichlet boundary condition (*ϕ_s_=* V). An insulating Neumann boundary condition was defined at the other sides of the cube 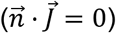.

The electrode-electrolyte interface at the surface of the implant was modeled using the Butler-Volmer equation [16], which expresses the current density as an antisymmetric function of the overpotential *η* at the interface (the voltage with respect to the equilibrium voltage),

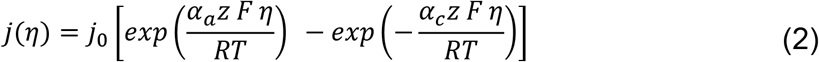

where *j_0_* is the exchange current density (in units of A/m^2^), *α_a_* and *α_c_* are the anodal and cathodal charge transfer coefficients (non-dimensional), *T* is the temperature (set at 310 K) and *R* and *F* are the universal gas constant and Faraday’s constant respectively.

The steady state solution of the problem described above was calculated using FEM in COMSOL. Unless stated otherwise, the exchange current density was set to *j_0_* = 10^−3^ X/m^2^ and the equilibrium voltage was set to −400 *mV,* which are values within the range of the reported values for Ti alloys in saline or biological media [17–19]. The charge transfer coefficients *α_a_* and *α_c_*, were arbitrarily set to 0.5, which is equivalent to assuming that the reactions at the interface involve the exchange of one electron. The impact of all these parameters is analyzed in the results section. Finally, the conductivity of the medium surrounding the implant was set to 1 S/m. This value was selected to be in the same order of magnitude of many biological tissues (e.g., cerebrospinal fluid, scalp or grey matter [20]).

### Simple electrical box model

The same geometry described in the previous section was simulated with a purely ohmic electrical model, i.e., one with constant conductivity in each region. The boundary conditions on the outer surfaces of the cube were kept the same and three different models of the implant were simulated:

▪ Perfect conductor, imposing a floating potential boundary condition with zero current (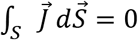 and *φ* ϕ *V* over all the surface, with *V* an unknown voltage)
▪ Perfect insulator, imposing a Neumann boundary condition 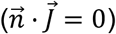.
▪ Usual boundary conditions at the interface of media with two different electrical conductivity that impose continuity of current perpendicular to the surface and of the tangential component of electric field (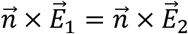 and 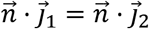)

As in the previous section, the electric potential distribution for the different models was calculated by finding the steady state solution of Laplace equation (2) with COMSOL.

### Realistic model of tES with skull plate

To obtain a 3-D representation of a skull plate, a picture of a circular plate taken from Rotenberg et al. [7] was binarized in MATLAB. Manual corrections of the binarized image were made using GIMP (https://www.gimp.org/, a free source image editing software) and the resulting image was copied into various slices (equivalent to a 0.7 mm thickness of the plate) to generate a voxel based volumetric representation of the plate. Then an iso2mesh [21] function was used to generate a tetrahedral mesh from this volumetric representation of the plate (see Figure 2a).

**Figure 2.**
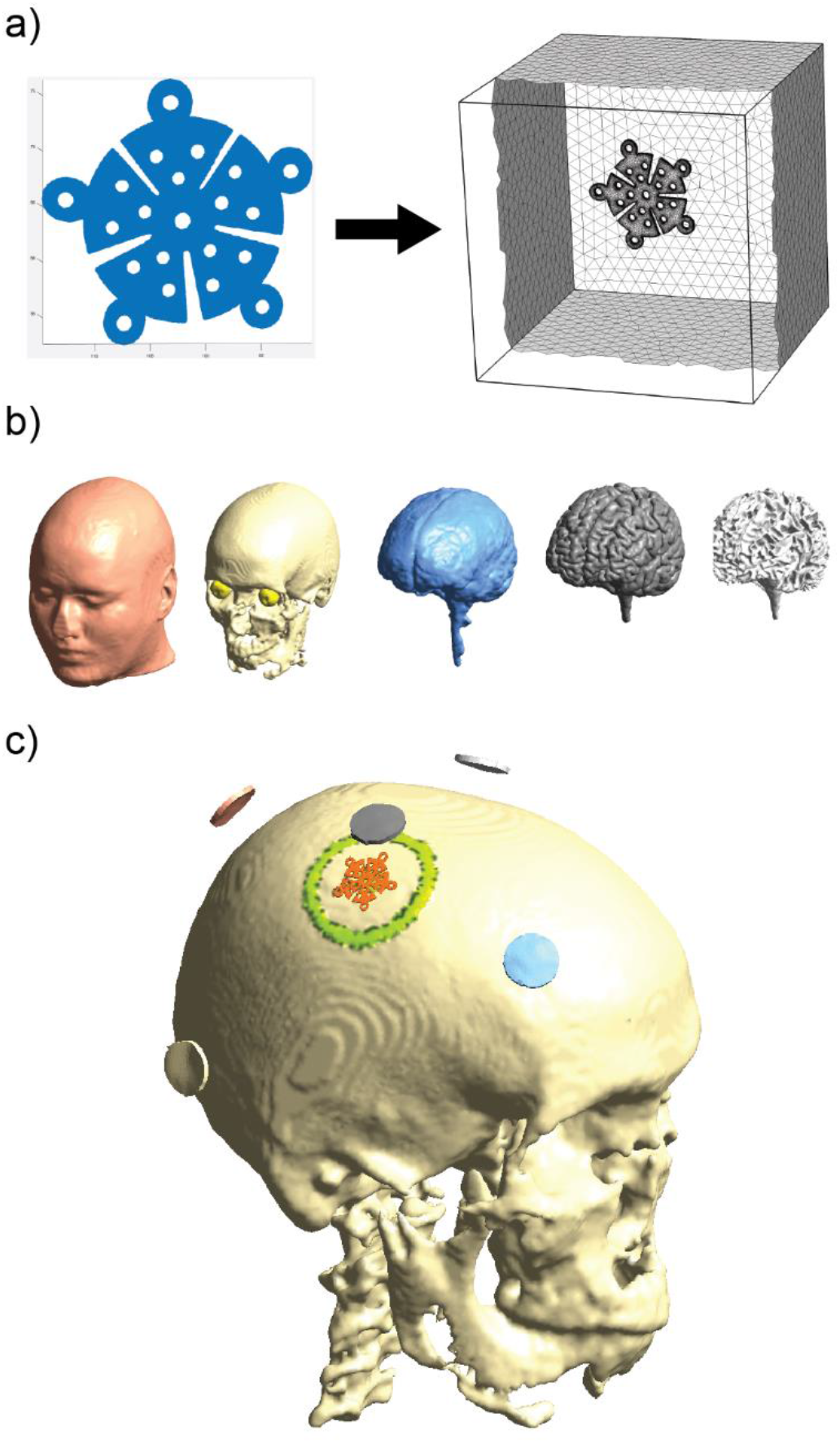
a) 2-D image of the skull plate used to generate a three-dimensional tetrahedral mesh of it embedded in a box. b) 5-layer head model used to simulate a tES treatment. c) Schematic of the electrode montaged simulated. A current of 2 mA was set on the C2 electrode (electrode above the plate) and a current of −0.5 mA on C1, F4, P8 and Pz electrodes. This electrode montage was selected to create a worst-case scenario by inducing a large current near the plate (orange) and the craniotomy (yellow).

To simulate the plate exposed to an external electric field, the plate mesh was imported into COMSOL. A 60 mm cube was created to surround the plate and the resulting geometry was meshed using COMSOL’s built in tools. The rest of the steps were the same as described in previous sections. Briefly, a homogeneous field was generated by imposing a voltage difference between opposite faces of the cube and the rest of the faces were considered insulators. The plate interface was modeled with the Butler-Volmer equation (2) using the same parameters described previously (parameters for titanium) and the electric potential distribution was calculated by finding the steady-state solution to Laplace equation.

To simulate a realistic tES treatment, we used the head model of the subject “Ernie” from SimNibs example dataset [22]. For this subject, a tetrahedral mesh is readily available with domains for scalp, skull, cerebrospinal fluid (CSF), grey matter (GM), white matter (WM) and eyes (see Figure 2b). A skull hole representing a craniotomy and a skull plate were added to this healthy head model through local mesh refinement using iso2mesh [21]. Briefly, the process consisted of, first, generating a tetrahedral mesh for 3 concentric cylinders with radii of 5, 20 and 24 mm to create the craniotomy. The inner cylinder represents a central hole and the two outer cylinders delineate a circular bone flap with a 4 mm thick gap. Second, the nodes of the plate mesh were translated and rotated to locate it at the craniotomy and aligned perpendicular to the skull. Finally, the nodes of these two meshes were added to the healthy head mesh. The resulting mesh can be seen in Figure 2c.

The head mesh with the craniotomy and the skull plate was used to simulate a tES treatment. The electrode montage consisted of 5 round, 1 cm radius, electrodes (NG PiStim, Neuroelectrics Inc.) placed over the C1, C2, F4, P8 and Pz positions of the 10/20 electroencephalography system. A current of 2mA was set on C2 and a current of −0.5 mA was set on the rest of the electrodes. Note that this montage was not meant to target any specific brain area but to simulate a worst-case scenario where the plate is located right underneath an electrode with a strong current and, thus, strongly interferes with the field distribution generated at the brain (see Figure 2c).

The electric field distribution generated by the montage described above was calculated using SimNibs 3.2. The tissues were modeled as isotropic with conductivity values of 0.33 S/m for the scalp and the eyes, 0.008 S/m for the skull, 1.79 S/m for the CSF, 0.40 S/m for the GM and 0.15 S/m or the WM [23–25]. The craniotomy’s burr-hole was modeled with the same conductivity as the CSF which corresponds to an acute state and constitutes the worst-case scenario [2]. The electrodes were represented by simply adding a geometrical representation of the gel underneath them (3 mm thickness cylinders with the same radius of the electrodes) with a conductivity of 4 S/m. Finally, the model was solved for two different plate conductivities, 10^−6^ S/m and 10^6^ S/m, to approximate the perfect insulator and perfect conductor situations respectively.

## Results

### Electrochemical and electrical models

Figure 3 a and b display the electric field magnitude and the electric field lines obtained in a cut plane of the geometry when modeling the implant as a perfect conductor or as a perfect insulator respectively. When the implant is modeled as a perfect conductor, the electric field lines tend to collapse at its surface, perpendicularly to it (as it should, the electric field is normal to the surface of a perfect conductor). This causes a local increase in the field magnitude near the surface regions that are perpendicular to the field and a local decrease near the surface regions that are parallel to it. In contrast, if the implant is modeled as a perfect insulator, the electric field lines pass around the surface circling it (the electric field is tangential at the surface of a perfect insulator). This causes the opposite effect in the field magnitude near the surface: a local decrease near the surface regions perpendicular to the field and a local increase near the regions parallel to it.

**Figure 3:**
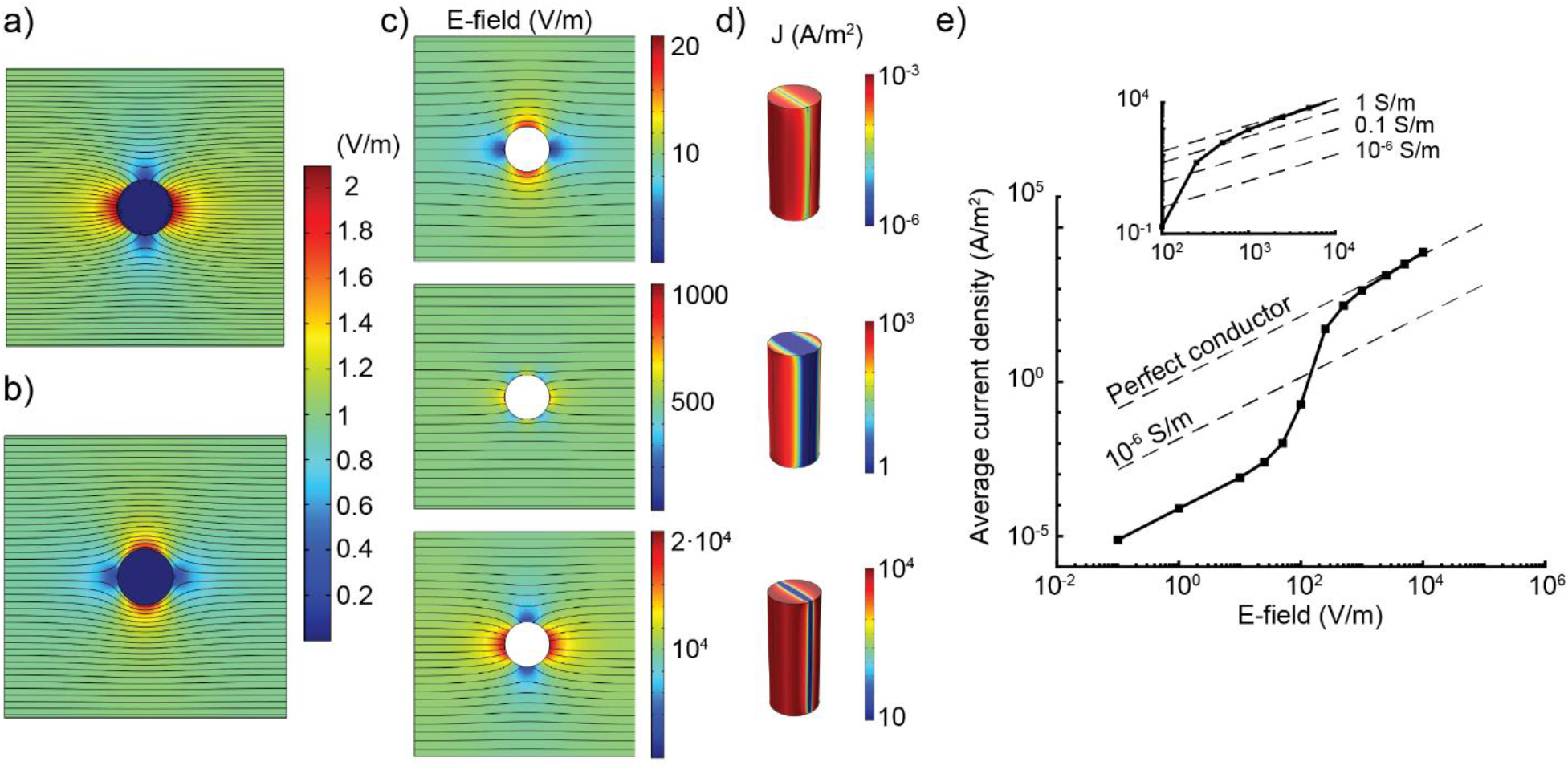
Simple box model. a) Electric field magnitude distribution and electric field lines in a plane perpendicular to the implant axis obtained with the electrical model when the implant is modeled as a perfect conductor. b) Same as a) when the implant is modeled as a perfect insulator. c) Electric field magnitude distribution obtained with the electrochemical model in a plane perpendicular to the implant axis for external electric field magnitudes of 10, 500 and 10000 V/m. d) Current density at the implant surface for the same electric fields. e) Average current density flowing through the implant surface as a function of the external electric field. The squares and the solid line provide the results obtained with the electrochemical model and the dashed lines show the results obtained with the electrical model when the implant is modeled as a perfect conductor or with different conductivity values.

We generated analogous plots using the electrochemical model. Figure 3c displays the field magnitude distribution for three different values of the applied external electric field. The results show that the spatial distribution of the field changes depending on its magnitude. When an external field of 10 V/m is applied, the field distribution resembles that obtained when modeling the implant as an insulator. On the other extreme, for a field of 10 kV/m we obtained a spatial distribution that is analogous to that obtained modeling the implant as a perfect conductor. Finally, between these two values, for a field of 500 V/m, a mixed pattern was obtained.

Figure 3d shows the current density through the implant surface for the same electric field values. When the implant is exposed to field magnitudes of 10 V/m or 10 kV/m, the current density has a value within the same order of magnitude over most of the surface. In contrast, when the implant is exposed to 500 V/m, two distinct regions with several orders of magnitude difference in the current density and a sharp transition between them can be observed. This means that, at this field strength, part of the surface allows charge transfer (acting as highly conductive) and the rest does not allow it (acting as highly insulating). This explains the pattern in the field distribution near the implant shown in Figure 3d.

To further analyze this behavior, we computed the mean current density through the implant surface as a function of the external electric field. This was done using COMSOL built-in functions. The results are presented in Figure 3e. For the sake of comparison, the results obtained with the electrical model using different properties for the implant are also presented. At low electric fields (less than 100 V/m), the electrochemical model shows an ohmic behavior, with a linear relationship between the electric field and the current density. The proportionality constant between them (i.e., the conductivity) is very low. Indeed, the results within this range of fields can be replicated with a purely electrical model defining a very low conductivity (below 10^−6^ S/m) at the implant making it act in practice as a perfect insulator.

In the range of fields between 100-2000 V/m, there is a transition region where the linear relationship between the field and the current density is lost. As shown by Figure 3d, within this range of fields, parts of the surface act as an insulator and others act as a conductor. Thus, as the field increases, a larger portion of the surface allows charge transfer causing a non-linear dependence between the field and the current density. It is worth noting that, except for the highest end of fields, within this transition region the electrical conductivity that would produce the same results is far from that of a good conductor. In fact, only at fields above 1000 V the equivalent conductivity is larger than that of the surrounding medium.

Finally, at electric fields above 2000 V/m the electrochemical model displays again an ohmic behavior. However, unlike the behavior at low fields, the equivalent conductivity is much higher. In fact, the results are identical to those obtained when modeling the implant as a perfect conductor in the electrical model.

### Influence of the parameters in the electrochemical model

To analyze the impact of the parameters used in the Butler-Volmer equation, we computed the average current density through the implant’s surface for different sets of parameters. Figure 4a shows the results obtained with different values of the exchange current density with other parameters fixed. In all cases, as described in the previous section, we observe an ohmic behavior at low electric fields followed by a transition region and another ohmic region at high electric fields.

**Figure 4:**
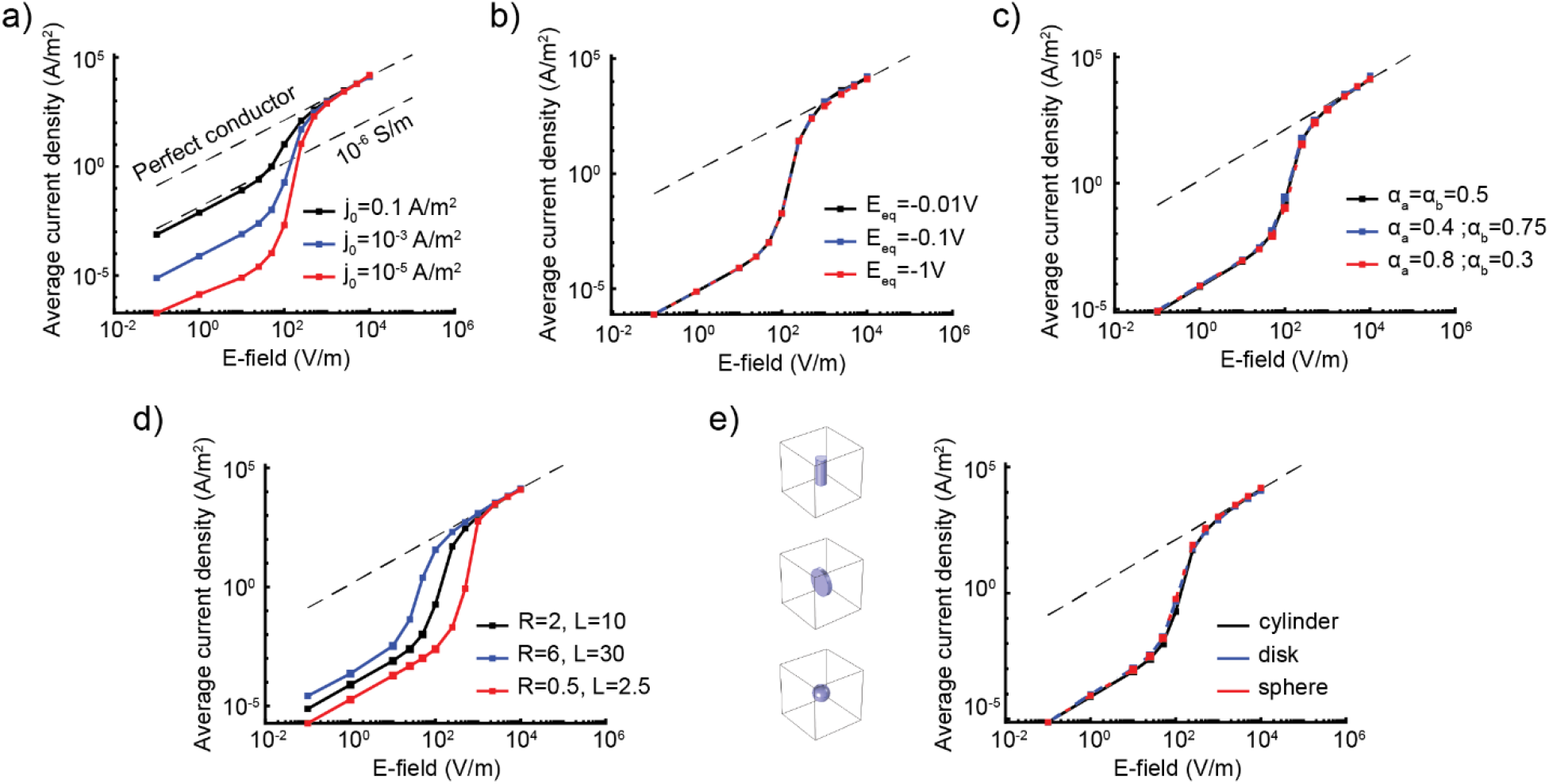
Average current density flowing through the implant surface as a function of the external electric field for: a) different values of the exchange current density, *j_0_*, (*α*_a_ ϕ *α*_c_ ϕ 0.5 and *E_eq_* ϕ −0.5 *mV*) b) different values of the equilibrium potential, *E_eq_* (*j*_0_ ϕ 10^−4^ *A/m^2^* and *α_a_* ϕ *α_c_* ϕ 0.5), c) different values of the charge transfer coefficients *α_a_* and *α_c_* (*j_0_* ϕ 10^−3^ *A/m^2^* and *E_eq_* ϕ −0.5 *mV*), d) different dimensions of the cylindrical implant keeping the ratio between its radius (R, in mm) and its length (L, in mm) constant and e) different shapes of the implant with the same volume. The dashed lines show the results obtained with the electrical model when the implant is modeled as a perfect conductor or with a specific conductivity value.

The lower and upper bounds of the transition region are not affected by the exchange current density. Similarly, for high electric fields all models tend to the same results (perfect conductor) regardless of the exchange current density. However, for low electric fields, although preserving a linear relationship, the proportionality constant is different depending on the exchange current density. In other words, the behavior is qualitatively the same regardless of the exchange current density, but, at low electric fields, the equivalent conductivity of the implant increases as the exchange current density increases.

The effects of changing the equilibrium potential of the implant are displayed in Figure 4b. In this case, the results are identical for all values tested, meaning that the equilibrium potential does not have a significant impact on the overall charge transfer at the interface when the implant is exposed to an electric field. Figure 4c shows the same results but for different values of the charge transfer coefficients. Similarly, these parameters do not have a significant impact on the qualitative behavior of the system. Although the charge transfer at the interface will be affected by the charge transfer coefficient (especially local differences are expected), the upper and lower bounds of the transition region are the same and the overall current density is very similar in all cases.

We also analyzed the influence of the geometry of the implant. Figure 4d shows the results obtained when the cylinder dimensions are increased or decreased keeping the same ratio between its radius and length. The same behavior is observed independently of the size. However, the transition region is displaced depending on the implant’s size. Whereas increasing the implant’s size moves the transition region towards lower fields, decreasing it has the opposite effect. In fact, there is an order of magnitude difference between the field at which the largest and the smallest cylinders simulated behave as conductors.

Finally, we generated the same results for different implant shapes having approximately the same volume. Namely, we simulated two additional shapes: a disk with a radius of 5 mm and a thickness of 1.5 mm and a sphere with a radius of 3.1 mm. The obtained results are displayed in Figure 4e. Unlike the size, the shape has a small impact on the results as only a slight difference is observed for the different shapes of the implant.

### Test case: tES treatment in a subject with a skull plate

To illustrate the importance of the correct choice of the electrical parameters, we simulated a tES treatment in a subject with a craniotomy and a skull plate. First, we simulated the skull plate alone, surrounded by a medium with a conductivity of 1 S/m and exposed to a homogeneous electric field, using the electrochemical model. Figure 5a displays the average current density through the plate surface as a function of the electric field. The results are very similar to those reported in the previous sections. Namely, the plate acts as an insulator until the external field reaches values in the order of thousands of V/m.

**Figure 5:**
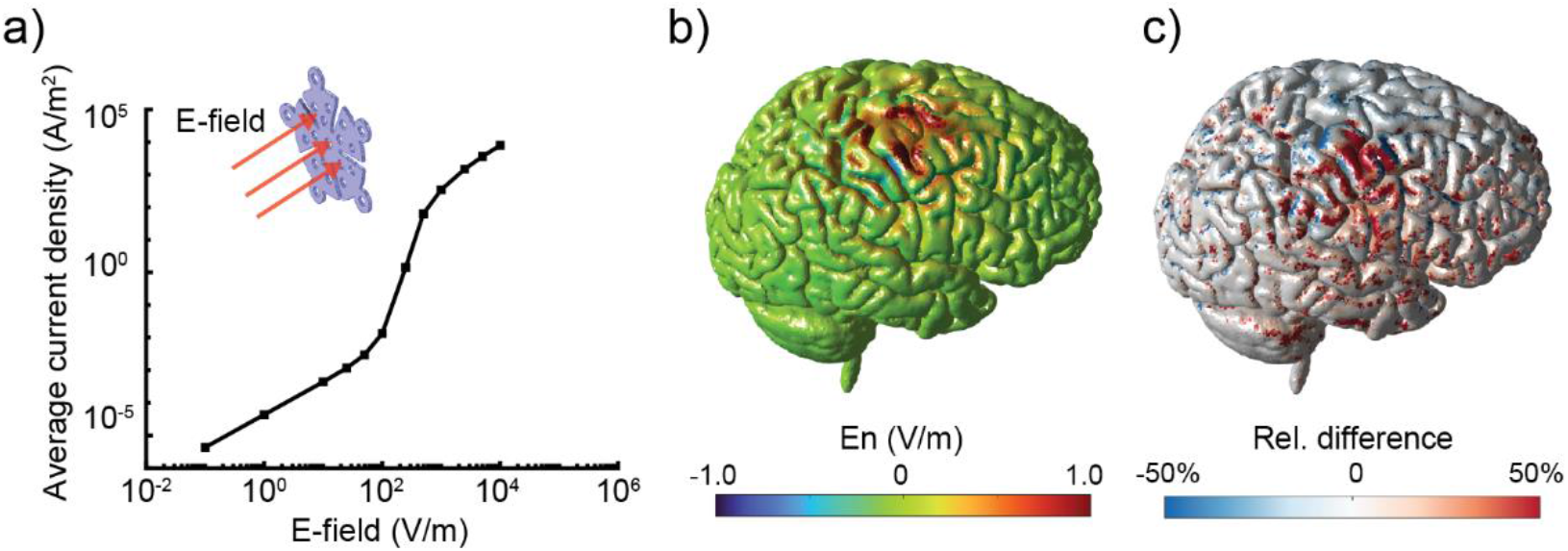
a) Average current density flowing through the surface of the skull plate as a function of the external electric field magnitude when it is exposed to a homogeneous electric field perpendicular to it. b) Normal component of the electric field at the cortex obtained when applying the tES treatment depicted in Figure 2c defining a conductivity of 10^−6^ S/m at the skull plate. c) Relative difference in the normal component of the electric field (in %) at the cortex between the models with a conductivity of 10^−6^ S/m and 10^6^ S/m assigned to the plate.

Then, we simulated the tES treatment with an electrode montage that was chosen to target the skull plate instead of a specific brain region (see methods section). The normal component of the electric field at the cortex was computed (see Figure 5b) by assigning a low conductivity (10^−6^ S/m) and a high conductivity (10^6^ S/m) to the skull plate. Figure 5c shows the relative difference between the two models. Errors above 50% are found in large regions of the brain underneath the plate.

## Discussion

Our results show that, despite the complex electrode-electrolyte interface phenomena, for most purposes the surface of an implant can be modeled with relatively simple approximations (perfect conductor or perfect insulator) and using a purely ohmic electrical model. In other words, generally the electric field distribution can be calculated accurately while ignoring the electrochemical nature of the problem with the correct choice for the boundary conditions used to define the implant’s surface.

The exponential nature of the Butler-Volmer equation that describes the electrode-electrolyte interface causes metals to act as perfect insulators when exposed to low electric fields and as perfect conductors when exposed to high electric fields. Between these two states there is a transition region in which the equivalent electric conductivity increases abruptly going from highly insulating to highly conductive with only an order of magnitude increase in the field. It is important to note that the conductivity increase within this range is associated to a larger portion of the surface allowing charge transfer and acting as a good conductor This means that it is not possible to approximate the situation with an adapted constant conductivity of the implant.

The parameters of the Butler-Volmer model depend on the materials involved (metal and electrolyte) and the reactions taking place at the interface. The equilibrium potential usually has values within ±3 V [26] and the exchange current density can vary within a very wide range – from 1 to 10^−9^ A/m^2^ [16]. The charge transfer rates depend on the number of electrons that are exchanged throughout the interface reactions. It is very common to assume that one electrode is exchanged in the anodal and cathodal reactions (with an associated transfer rate constant of 0.5). These parameters are very relevant when modeling corrosion kinetics or electrochemical cells, but fortunately, for our purposes they don’t have a significant impact. None of the parameters affects significantly the upper and lower bounds of the transition region between the insulating and the conductive behavior or the implant. Only the charge transfer coefficients can have a mild effect on them. This is indeed readily explained by the model equation (2) and its exponential nature. The exponent depends linearly on the voltage at the interface, making it the determining factor for this transition from insulator to conductor. Since the voltage depends mostly on the external field and implant geometry, the transition region bounds are largely independent of the parameters. Nonetheless, although not modifying the bounds of the transition region, the exchange current density determines the equivalent conductivity at the insulator regime. Yet, even at the upper limit of the values that are typically reported for this parameter, the equivalent conductivity is very low (below 10^−6^ S/m).

Unlike the Butler-Volmer equation parameters, (i.e., the type of metal the implant is made of and surrounding media), the geometry of the implant has an impact on the qualitative behavior at the interface for different electric fields. This can be expected since geometry has a great impact on the voltage induced at the interface for a certain field. Thus, the transition from insulator to conductor will occur at different values of the field depending on interface location, shape and size the implant has. Indeed, as shown by our results, changing implant volume but preserving shape displaces the transition region significantly, while changing implant shape with constant volume does not. Thus, the electric field at which the implant will change its behavior from insulator to conductor mostly depends on its size.

We simulated a tES treatment applied to a subject with a skull plate as a test case to illustrate the relevance of the analysis presented in this study. We first modeled the plate exposed to a homogeneous external electric field using the electrochemical model. Based on these results (Figure 5a), we can approximate the plate as a perfect insulator for fields up to a few hundreds of V/m and only at fields above 2000 V/m the plate acts as a perfect conductor. This results, combined with the peak values of the electric field in the head reported for a tES treatment [27], warrant modeling the plate as an insulator. Modeling the plate as a perfect conductor can result in large errors on the dosimetry assessment. Indeed, the estimates of normal electric field at the cortical surface obtained with high or low conductivity to the plate display a significant difference in large regions of the brain in our simulations.

It is important to note that the analysis presented here applies for any electrical stimulation technique operating in the quasi-static approximation regime, i.e., with frequencies in the low kHz range and below, where Faradaic, as opposed to capacitive, currents dominate. Among brain stimulation techniques, this includes most tES modalities, TMS and electroconvulsive therapy (ECT). According to numerical models, the maximum electric field values over the entire head in tES are around 30 V/m [27] and in TMS around 300 V/m [28]. Our results indicate that, as a rule of thumb, any implanted metal in the head can be electrically modeled as a perfect insulator in the context of tES or TMS. In ECT the electric field reaches higher values (above 4500 V/m at the scalp [29]). In this case, the choice of the correct electrical model for an implant may not be straightforward as it will depend on its geometry and how strong the electric field is expected to be in its location.

A similar reasoning can be applied to any technique that is based on the delivery of electric currents, such as functional electrical stimulation or electroporation-based treatments. It may also be relevant when modeling implanted electrodes for voltage measurements or non-active contacts of a stimulating electrode. In these cases, a low interface voltage is generally expected at non-active contacts, especially in measuring electrodes. Thus, according to our analysis, electrodes in those cases should usually be modeled as insulators. In fact, this is aligned with a prior study that compared a detailed impedance model with an insulator boundary condition as means to represent a microelectrode array used to record an action potential [30]. The authors of that study concluded that, the electrode-electrolyte interface could simply be modeled with an insulating boundary condition without the need to implement more complex impedance models.

## Conclusions

Although implanted metals have a high nominal electrical conductivity, modeling as such is incorrect in important cases, and especially in tES and TMS. The nature of the electric currents is different between metals and biological tissues (electronic vs ionic). Consequently, current can only flow through the surface of an implant by means of electrochemical reactions which in turn need a minimum voltage gradient to occur. As illustrated by our results, this causes the charge transfer between a metal implant and the surrounding tissue to be insignificant unless the implant is exposed to a high enough electric field. Thus, metal implants act as perfect insulators unless a sufficient voltage is induced at their interface. Only when implants are exposed to a large electric field a sufficient voltage is induced at the interface allowing charge transfer. When this happens, the high nominal conductivity of the implant is reflected in the current distribution resembling the behavior of a perfect conductor. The transition between these two states is abrupt enough to assume that implants can generally be modeled either as highly insulating or as highly conductive, except in a narrow transition region which may be important in ECT. Thus, with the correct model choice, the electrochemical nature of the problem can be ignored. However, as we showed with a test case, the incorrect model choice can lead to large errors in the estimation of the electric field distribution.

## Acknowledgements

This work has received funding from the European Research Council (ERC) under the European Union’s Horizon 2020 research and innovation programme (grant agreement No 855109) and from FET under the European Union’s Horizon 2020 research and innovation programme (grant agreement No 101017716)

